# The nutritional content of anthropogenic resources affects wildlife disease dynamics

**DOI:** 10.1101/2025.04.10.648226

**Authors:** Erin L. Sauer, Jessica L. Hite, Sarah E. DuRant

## Abstract

Wildlife have become increasingly reliant on human-supplemented food, affecting interactions between individuals and subsequently disease transmission. Much of the research investigating the impact of human-supplemented food has focused on host behavioral changes and food availability. However, the nutritional quality of food can significantly impact disease transmission depending on whether it boosts or hinders immunity. Few studies have investigated links between nutrition, immunity, and pathogen transmission, which weakens efforts to guide and manage infectious disease at the wildlife-human interface. We investigate these interactions using a data-driven transmission model that explicitly incorporates feedback between nutrition, immunity, and pathogen transmission in the bacterial pathogen, *Mycoplasma gallisepticum* and its avian hosts which are heavily dependent on supplemental food sources. We also examine the nutritional content and sales frequency of one of, if not the, most common source of anthropogenic food supplementation, bird feeder food. We find that food with lower nutritional quality is purchased for wild bird supplementation far more frequently than food with high nutritional quality. Further, our transmission model demonstrates that these lower quality food sources result in large epidemics that cause destabilizing population declines. Our results suggest that nutritional quality plays an important role in disease transmission and provide insight into the ecological and immunological consequences of human behavior on infectious disease in wildlife.

## Introduction

Human activity has altered the availability, quality, and distribution of resources for wildlife. This is particularly evident in urban and suburban systems, where wildlife are increasingly adapting to environments that provide abundant anthropogenic food sources, such as discarded food, pet food, bird feeders, and urban gardens (Schell et al., 2021). As a result, wildlife in human-dominated areas are becoming more reliant on these human-provided resources (Robb et al., 2008). This shift can disrupt natural behaviors and hierarchies, contributing to an increased risk of disease transmission, as animals come into closer contact with humans and each other in urban spaces (Becker et al., 2015; Combs et al., 2022). Reliance on anthropogenic food sources can also alter the nutritional quality of food, limiting caloric availability and/or the availability of specific nutrients that enable hosts to effectively combat infectious disease (Murray et al., 2016). Given these substantial shifts in resource availability and their potential impacts on wildlife and human health, evidence-based policy changes will be critical for conservation and to protect the health of domestic animals and humans. Yet, we are far from being able to make such recommendations in most systems – in part because pathogen transmission depends on multiple non-linear interactions, especially with diet-immunity links, that are difficult to disentangle. To date, most studies have focused on subsets of these links and on food quantity rather than food quality. Data-driven mathematical models can provide significant advances by linking these siloed data into an integrative framework.

The mechanisms by which anthropogenic food sources alter disease dynamics in wildlife are complex and there is evidence for both positive and negative effects on wildlife health and disease risk (Becker et al., 2015; Murray et al., 2016). For instance, increased host density and contact rates at anthropogenic food sources can elevate transmission within and between species, a phenomenon that has been well-documented in studies of wildlife (reviewed by Becker et al., 2015; Murray et al., 2019). Aggregation at food sources can also result in immunosuppression caused by increased aggressive encounters and chronic stress (Murray et al., 2016; C. A. D. Semeniuk et al., 2009; C. Semeniuk & Rothley, 2008). Additionally, the quality and availability of food can have immunological consequences for hosts that alter disease transmission (Hite et al., 2020; Murray et al., 2016). For example, high bird feeder availability resulted in increased innate immunity of black-capped chickadees (*Poecile atricapillus*) while human food heavy diets (bread) in white ibis (*Eudocimus albus*) resulted in decreased innate immunity (Cornelius Ruhs et al., 2019; Cummings et al., 2020). These effects on immune function can directly influence pathogen transmission, as greater pathology or pathogen growth may increase infection likelihood and intensity and increase recovery times (Cornet et al., 2014; Hite & Cressler, 2019; Murray et al., 2016). Together, these studies highlight how human-supplemented food can reshape disease transmission by influencing host density, behavior, and nutritional status.

Discerning the roles of aggregation from dietary mediated changes on immune function in driving pathogen transmission can be challenging (Altizer et al., 2018). Most attempts have focused on food availability (Strandin et al., 2018). Many studies have shown wildlife in areas with access to anthropogenic sources of food have improved innate immune function (Cornelius Ruhs et al., 2019; French et al., 2022; Hwang et al., 2018; Strandin et al., 2018), though prioritization of innate immunity could be a response to increased pathogen exposure and/or low-quality food (Becker et al., 2018). While abundant resources can be beneficial, the quality of anthropogenic foods can vary greatly and create trade-offs for wildlife (Brown & Cooper, 2006; Jones et al., 2014; Murray et al., 2016; Stewart et al., 2016; Strandin et al., 2018). For example, one study found that urban white ibis (*Eudocimus albus*) had fewer ectoparasites due to reduced effort finding food but also had reduced body condition due to the poor quality of their diets (Murray et al., 2018). Similarly, a study on coyotes (*Canis latrans*) found that while urban coyotes have greater access to a diverse diet, low-protein anthropogenic resources increased risk for disease and human conflict (Murray et al., 2015). Thus, assessing the quality of anthropogenic food and its effect on immune function presents an additional hurdle, as studies need to account for differences in both food availability and nutritional quality.

The quality of anthropogenic food can have significant effects on wildlife health and immune function, influencing disease dynamics in complex ways. Contaminants such as mycotoxins, produced by fungi in poorly stored or processed food, can suppress immune responses, making wildlife more susceptible to infections (Murray et al., 2016). Additionally, anthropogenic foods can alter host microbiomes, which in turn can impact immune function, though the precise effects vary depending on the species and the specific food source (Fleischer et al., 2024; Ingala et al., 2019; Knutie, 2020; Love et al., 2024; Viquez-R et al., 2024). Furthermore, the balance of macro- and micronutrients in food is essential for immune cell production and overall health. For instance, dietary proteins are crucial for immune function, and insufficient intake can impair a host’s ability to respond to infections (Taylor et al., 2013). Prior research on canaries has shown that dietary macronutrients play a critical role in immune function and tolerance to bacterial and parasitic infections (Cornet et al., 2014; Perrine et al., 2025; Sauer et al., 2024). These factors demonstrate how dietary quality, beyond just food availability, plays a pivotal role in determining disease risk and immune responses in wildlife. However, studies linking the effects of dietary quality on immune function to pathogen transmission are limited.

Here we examine the nutritional quality of food commonly sold for wild birds and link that to the potential for dietary quality to drive transmission through its effect on host immune function in the widespread avian bacterial pathogen *Mycoplasma gallisepticum* (hereafter MG). MG is a conservation concern for wild birds, causing large declines in house finch (*Haemorhous mexicanus*) populations and outbreaks in other bird species after its emergence in the mid-1990’s (Dhondt et al., 2014; Hochachka & Dhondt, 2000; Kollias et al., 2004). MG is also an economic problem for poultry farming, costing the poultry industry hundreds of millions of dollars annually in the US alone and contributing heavily to the use of antibiotics in livestock (Ley & Yoder Jr, 2008; Yadav et al., 2022). MG is transmitted via fomites and direct contact with infected birds, with bird feeders playing a key role in driving transmission within the wild system (Adelman et al., 2015; Dhondt et al., 2007; Hochachka et al., 2021; Hochachka & Dhondt, 2000; Moyers et al., 2018). Further, the introduced eastern house finch population is primarily found in urban and suburban areas, where supplemental food from bird feeders constitutes a significant part of their diet (Badyaev et al., 2020). Because of the crucial role bird feeders play in this system, the quality of food birds receive from them is likely to influence host immune function and thus affect disease dynamics (Perrine et al., 2025; Sauer et al., 2024).

Few studies in any wildlife system examine the potential for nutritional quality, as opposed to food quantity or availability, to affect disease transmission (Altizer et al., 2018; Murray et al., 2016). This knowledge gap is likely due to these types of studies being logistically difficult because they often require experimental feeding and infection studies on wildlife in captivity. Because house finches rely on bird feeders for food and the contribution feeders already play in transmission as fomites, MG is an ideal system for answering questions regarding how the nutritional quality of food offered at bird feeders contributes to disease severity and transmission in the system. To determine the potential for dietary quality to drive transmission in MG, we developed a multistate transmission model parameterized with exposure-controlled pathological data from domestic canaries fed diets of varying nutritional qualities (*Serinus canaria domestica*) and data from house finch studies. Canaries exhibit similar pathology and susceptibility to MG as house finches while being better suited for laboratory conditions, making them an ideal model for the system (Hawley et al., 2011). Prior research on canaries demonstrates that dietary macronutrients play a critical role in immune function and tolerance to MG, yet how these effects scale-up to shape transmission dynamics remains unresolved (Perrine et al., 2025; Sauer et al., 2024). We also present an analysis of wild bird food sales data from a local bird store specializing in supplemental food for wild birds. In addition to guiding our study design, these real-world data help contextualize our results in the wild system and provide information that can help inform well-intentioned bird-enthusiasts. While numerous studies have examined various aspects of the multi-faceted challenge of supplemental feeding, few have directly connected human-supplemented bird food nutritional quality that reflects consumer data.

## Methods

### Bird food sales: data acquisition

We acquired bird food sales data (December 2023-November 2024) and nutritional information (% minimum crude protein and fat content) from a local store specializing in supplemental food for wild birds (Wild Birds Unlimited Nature Shop, Fayetteville, AR). Sales data does not include nectar. Because we were not able to acquire mass data for all bird food sales lines, sales frequency represents the number of units sold, regardless of volume or mass (e.g. a 10-lb bag of seed and 12-oz block of suet are counted the same).

### Dietary drivers of transmission: theoretical framework

We built a mechanistic model to investigate how nutritional quality impacts the epidemiological dynamics of MG. Specifically, we used simulations to compare the effects of high nutritional quality (high protein) and low nutritional (high lipid) quality on host population size (via virulence and immunopathology) and the size and duration of disease outbreaks. The compartmental S-I-R model divides the total population, *N(t)*, over time *t* into five states: susceptible, *S*, first-time infected, *I_1_*, recovered from first infection, *R_1_*, second-time infected, *I_2_*, and recovered from second infection, *R_2_*, states (Li & Muldowney, 1995). We conducted two simulations of our model, one for each dietary state (high-protein diet vs. high-lipid diet). The model assumes homogenous mixing of adult individuals, a density-dependent birth rate, and that individuals who recover for a second time move to the *R_2_* state where they are no longer susceptible (Hosseini et al., 2004). We end susceptibility after second recovery for multiple reasons: wild birds are unlikely to experience more than two infections due to life span and multiyear epidemic cycling (Hosseini et al., 2004), the contribution of these birds to transmission would be inconsequential due to improved immunity (Dhondt et al., 2012, 2017; Sudnick et al., 2025), and the model is not designed to accurately represent multiyear epidemic dynamics. Transmission dynamics are governed by the system of ordinary differential equations:

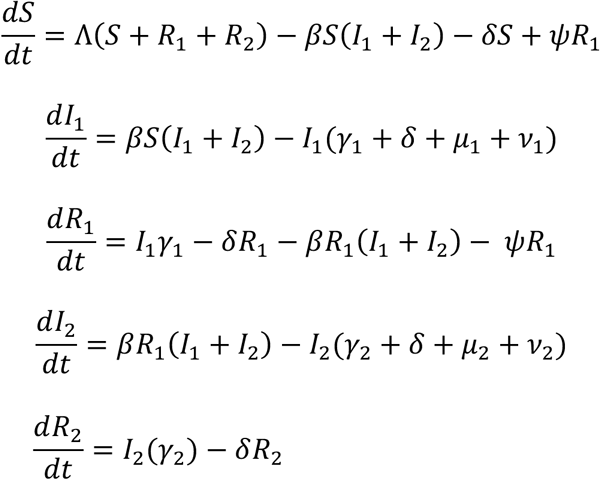

Simulations were conducted with R version 4.1.0 in R Studio using the *deSolve* package (RStudio Team, 2021; Soetaert et al., 2010).

We parameterized the model with a combination of data from existing studies and our recent infection assay. This experiment exposed individually isolated domestic canaries (*Serinus canaria domestica*) acclimated to either a high-protein (4:1 protein to lipid ratio) or high-lipid (1:4 P:L ratio) dietary regime to identical doses of MG (5.00x10^7^ CCU/mL; isolate VA1994; E. Tulman, University of Connecticut) then recorded disease outcomes including recovery time and immunopathology (Perrine et al., 2025). Recovery rate (*γ_1,2_*) and the additional mortality from infection-mediated inflammation *(µ_1,2_*) for each diet were estimated from the experimental infections in Perrine et al. (2025). Recovery rate from the first infection *(γ_1_*) was calculated as the inverse of the median number of days from exposure until recovery from symptomatic disease (i.e. eye score = 0). Recovery rate from the second infection was *γ_2 =_ γ_1_a* where *a* = 1.7, this value was based on the average fold change in recovery time from first to second infection in canaries (Sudnick et al., 2025). The inflammation-induced additional mortality rate *(µ_1,_ µ_2_*) was calculated as the daily probability of death following infection in isolation from other disease-induced causes of mortality (Cressler & Adelman, 2024). For individuals experiencing their first infection, the mortality rate was estimated using the following formula: *µ_1_* = −ln (1 − *N_d_*/*N*)/*D* where *N_d_/N* represents the proportion of individuals that died during the experiment, and *D* is the duration of the experiment in days. This approach accounts for the observed mortality over time and converts it into a daily rate. Taking the natural logarithm of this fraction and dividing by *D* gives the rate at which individuals die per day after exposure. The negative sign ensures the mortality rate is positive because ln(*x*) is negative when 0 < *x* < 1. Because partial immunity is expected to reduce mortality upon reinfection, we assumed that individuals experiencing a second infection had a lower risk of death. Specifically, the mortality rate for second infections (*µ_2_*) was set as *µ_2_* = *µ_1_*/2, reflecting a 50% reduction in mortality rate for second infections. This assumption aligns with previous studies indicating that prior exposure to MG can confer some protective effects against severe disease outcomes (Fleming-Davies et al., 2018; Sudnick et al., 2025).

The other parameters used in the model were either taken directly from or modified from studies of MG in house finches. The density-dependent reproductive rate function was taken directly from Cressler & Adelman (2024), using the same value for Λ and assumption that only susceptible (previously uninfected and recovered) individuals are reproducing. We assume that partial immunity of recovered individuals wanes at a rate of *ψ* and those individuals return to the susceptible state (Fleming-Davies et al., 2018). The rate of waning partial immunity (*ψ*) and infection induced mortality rate (expected mortality of infected birds in the wild due to things like increased predation) for first infection (ν_1_) was taken directly from Fleming-Davies et al. (2018). We assume that birds are more likely to survive a second infection due to partial immunity and thus decreased this mortality rate for second infections (ν_2_ = ν_1_/2). The transmission coefficient (*β*) for the bird fed a high quality (protein) diet was also taken directly from Fleming-Davies et al. (2018) and modified for lipid diet birds under the assumption that their transmission rate would be higher due to greater pathology and longer recovery times (β_protein_*2). We conducted a local sensitivity analysis to determine the robustness of our transmission coefficient estimates. For this analysis, we ran simulations of the previously described S-I-R model where β_lipid_ was calculated as either: β_protein_*2.5, β_protein_*2, or β_protein_*1.5. See Table 1 for full list of parameter definitions and their values.

**Table 1.** Epidemic model parameters, biological interpretations, and values for the protein diet population and the lipid diet population simulations.

| Parameter | Biological interpretation | Protein diet value | Lipid diet value |
| --- | --- | --- | --- |
| $\Lambda$ | Maximum density-independent birth rate | 0.025 | 0.025 |
| $\beta$ | Transmission rate | 0.00275 | 0.0055 |
| $\delta$ | Background host mortality rate | 0.01 | 0.01 |
| $\psi$ | Rate of waning partial immunity | 0.01 | 0.01 |
| $\gamma_1$ | Recovery rate from first infection | 0.048 | 0.033 |
| $\gamma_2$ | Recovery rate from second infection | 0.081 | 0.567 |
| $\mu_{1,2}$ | Inflammation-induced mortality rate | 0.003, 0.0015 | 0.009, 0.0045 |
| $\nu_{1,2}$ | Infection induced mortality rate | 0.25, 0.125 | 0.25, 0.125 |

The values for $\Lambda$ and $\delta$ and $\psi$ were taken directly from Cressler & Adelman (2024) and Fleming-Davies et al. (2018), respectively.

## Results

### Bird food sales: results

Sales frequency of supplemental food for wild birds in Fayetteville, AR, USA demonstrate that low protein to lipid (P:L) ratio bird foods are by far more frequently purchased than supplemental food with an even or high P:L (Figure 1).

**Figure 1.**
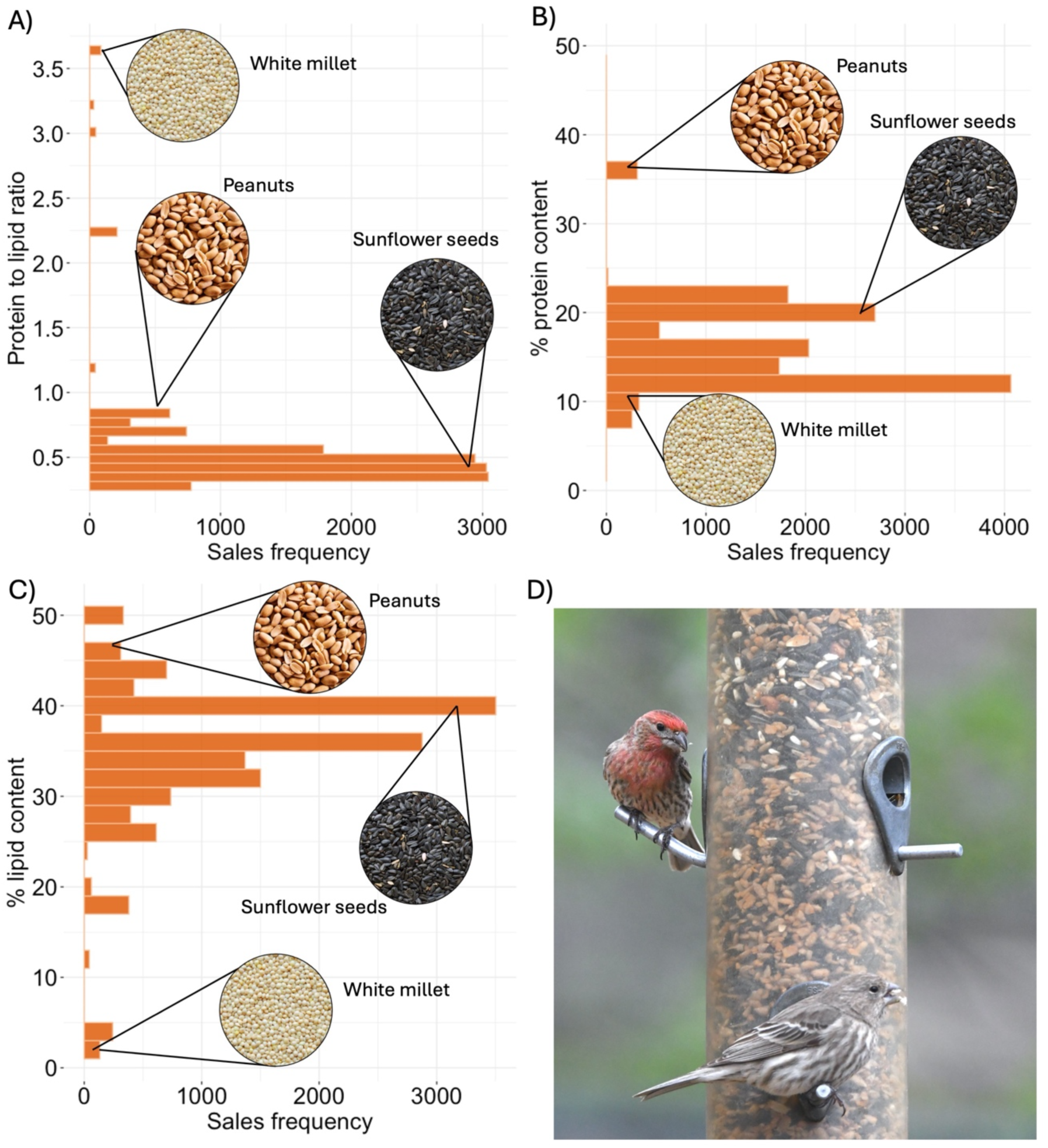
(A) Sales frequency of supplemental food for wild birds show that foods with lower nutritional content (lower protein to lipid (P:L) ratios and low protein content) far outsell food with higher nutritional value (high P:L ratio and high protein content). Ratios alone do not provide enough information on diet quality, and thus we examine sale frequency data on (B) protein and (C) lipid content per gram. Some high P:L foods, like white millet, actually have poor nutritional value with lower protein content than some more commonly purchased low P:L foods like sunflower seeds (B). To contrast, peanuts have a close to even P:L ratio but much higher protein content (B) and higher lipid content (C) than more frequently purchased sunflower seeds, making them a more nutritionally dense food option. (D) Two house finches on a bird feeder.

### Dietary drivers of transmission: results

Nutritional quality altered both the onset, duration, and severity of the epidemic. Low quality high-lipid diets resulted in large epidemics that began early and drove large population declines (Figure 1). The epidemic peaked within 25 days with a maximum of 34% infected birds, resulting in a 70% population reduction when the epidemic ended. In contrast, high-quality high-protein diets resulted in smaller epidemics with a lag in the onset of the outbreak and with minimal effects on host population densities. The epidemic took 60 days to peak with a maximum of 21% infected birds and no population reduction when the epidemic ended (Figure 2). Sensitivity analyses indicate the low-quality (lipid) diet always resulted in a greater proportion of infected birds and larger population declines than the high-quality protein diet, though transmission dynamics do become more similar the closer β_lipid_ is to β_protein_. These findings underscore the profound impact of nutritional quality on disease dynamics, with low-quality diets accelerating outbreaks, increasing infection severity, and driving significant population declines, while high-protein diets mitigate epidemic intensity and help protect host population stability.

**Figure 2.**
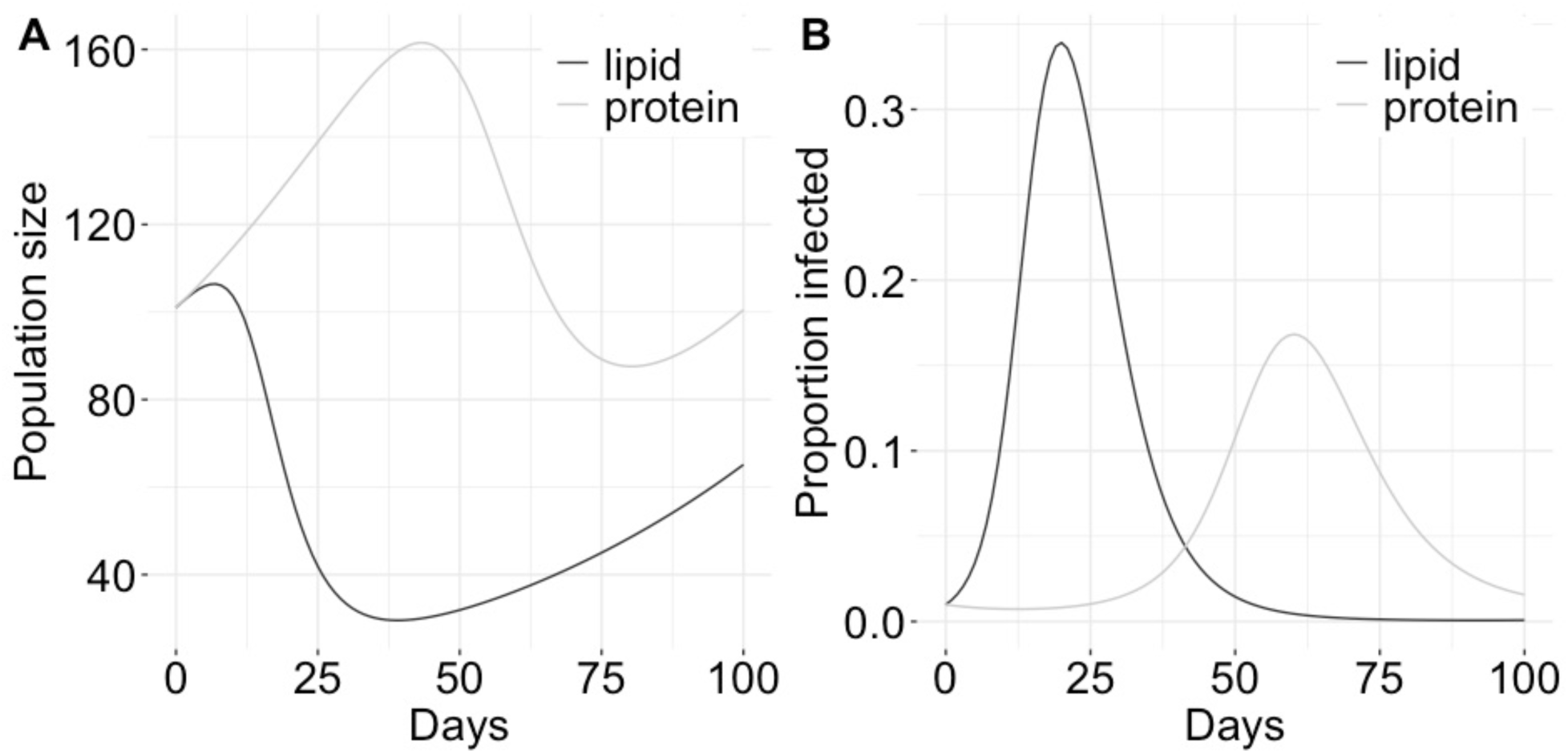
Results of S-I-R model simulations demonstrating the change in (A) total population size and (B) the proportion of the population that is infected over time by diet quality. The low-quality high-lipid diet (1:4 P:L ratio, dark-grey lines) resulted in larger epidemics with rapid onsets and greater population declines than the high-quality high-protein diet (4:1 P:L ratio, light-grey lines).

## Discussion

Here we aimed to examine how the nutritional quality of human-supplemented food affects disease transmission in a common avian host-pathogen system. We acquired nutritional content and sales frequency data from a retailer specializing in supplemental food for wild birds to guide and contextualize a data-driven transmission model that incorporates feedback between nutrition, immunity, and MG pathogen transmission. Bird feeders play an important role as both a primary food source for urban and suburban house finches and as points of pathogen accumulation and transmission. The bird food sales data showed that low-quality high-lipid foods are purchased far more frequently than high-quality high-protein foods. Our data-driven transmission model reveals that these low-quality high-lipid food sources result in large MG epidemics that cause destabilizing population declines. These results demonstrate the importance of using integrative approaches for determining the effects of human supplemented food on wildlife disease dynamics. While many studies have examined the impact of host aggregation and food quantity on transmission, few have incorporated nutritional quality and host immunity into transmission estimates and fewer still have connected human-supplemented bird food nutritional quality that reflects consumer data.

We find that lower quality foods with low P:L ratios and low protein content are more popular with people who provide supplemental food to birds than higher quality foods. There are several factors that may be influencing supplementers, including cost, retailer recommendations, and perceived preferences of the species people want to attract to their yards (Horn et al., 2014; Johansen et al., 2014). Results from Perrine et al. (2025) used here to parameterize the S-I-R model, show that birds fed a low P:L ratio diet had greater pathology, longer recovery times, and higher mortality than birds fed a high P:L ratio diet, possibly due to immunosuppression in the low P:L ratio birds (Sauer et al., 2024). Protein content is relatively low in the most purchased supplemental foods and even lower in other supplemented foods recommended for birds such as bakery scraps (Coogan et al., 2018; Horn et al., 2014). However, dietary proteins play an important role in immune function and protein deficient diets are associated with increased disease risk and decreased immune function in several invertebrates (Brunner et al., 2014; Cotter et al., 2019; Ponton et al., 2020; Povey et al., 2014) and vertebrates (Jahanian, 2009; Lochmiller et al., 1993; Taylor et al., 2013). Conversely, supplemental foods with high lipid content are very popular among supplementers. Lipids play a role in regulating inflammation processes, but high lipid diets can result in immune dysfunction in invertebrates (Adamo et al., 2010) and vertebrates (Sauer et al., 2024; Strandberg et al., 2009) as well as other negative consequences. For example, a study in blue tits found that supplemental feeding of lipids in the winter had negative effects on breeding (Plummer et al., 2013). Understanding the health consequences of shifts in the macronutrient availability in the urban environment due to supplementation and if wildlife are able to mitigate those shifts is an important area for future research (Coogan et al. 2018).

Our model demonstrated that a low-quality low P:L ratio diet led to large fast-paced epidemics and destabilizing population declines. These differences stem from slower recovery times and higher inflammation-induced mortality in the low-quality low-P:L diet simulations. Results from Sauer et al. (2024) on canaries suggest that prior to MG exposure, birds on the low-quality low-P:L diet have suppressed expression of immune process genes relative to birds on the high P:L diet. However, after exposure, canaries on the low P:L diet seem to overcompensate with high expression of immune and inflammation genes relative to high P:L diet birds resulting in greater immunopathology (i.e. greater conjunctival inflammation which lasts for a longer period and greater mortality) for the low-P:L birds. House finches are one of the most common visitors to bird feeders in the United States and Canada, making up 17% of all feeder visits according to one study and preferring low P:L ratio sunflower seeds (Horn et al., 2014; Johansen et al., 2014). The eastern house finch population in particular is almost exclusively found in urban and suburban habitats where supplemental food makes up a substantial portion of their diet (Badyaev et al., 2020). When MG first emerged in wild birds, feeder-dependent eastern house finch populations experienced substantial declines similar to those simulated in the low P:L ratio diet population of our S-I-R model (Hochachka & Dhondt, 2000). Our results provide evidence that the high P:L ratio supplemented diet of those populations may have contributed to immunopathology and subsequent declines in house finches.

Modern eastern house finch populations appear to have evolved tolerance to MG infection (Bonneaud et al., 2019; Henschen et al., 2023), though there is still substantial heterogeneity among individuals from co-evolved populations (Hawley et al., 2023; Henschen et al., 2023). Further, evidence suggests that modern isolates have co-evolved to become more virulent, resulting in a co-evolutionary arms race where immunopathology is necessary for transmission and tolerance results in selection for virulence (Bonneaud et al., 2019; Cressler & Adelman, 2024; Fleming-Davies et al., 2018; Hawley et al., 2023; Ruden & Adelman, 2021; Sauer et al., 2023). Thus, while the high-quality high P:L ratio populations had smaller epidemics, their tolerance to MG infection may have co-evolutionary consequences (Hite et al., 2020; Pike et al., 2019). Dietary nutrition, particularly in the feeder-dependent eastern populations, may be contributing to the heterogeneity in tolerance found in and among these populations. Further research into how the landscape of dietary quality in urban and suburban areas is contributing to heterogeneity in immunopathology during MG epidemics would help further our understanding of how the urban environment contributes to host-pathogen co-evolutionary dynamics (Coogan et al., 2018; Hite et al., 2020; Pike et al., 2019).

### Conclusions

The supplemental feeding of wild birds by humans is happening on an enormous scale and is certainly affecting the ecology and evolution of wild birds (Bosse et al., 2017; Robb et al., 2008; Shutt & Lees, 2021) and, in some instances, the non-avian species with which they interact (Orros & Fellowes, 2012; Shutt & Lees, 2021). The role supplemental feeding plays in disease is a complex multifaceted issue. Bird feeders can be hotspots for transmission (reviewed in Becker et al. 2015) and the nutritional consequences of supplemented food on immune function are varied. However, there is evidence that providing wild birds with supplemental food can increase the overall health of birds (Wilcoxen et al., 2015) and provides people with happiness and a feeling of being connected to nature (Cox & Gaston, 2016), which likely contributes to positive opinions about conservation (Dayer et al., 2019). People who provide birds with supplemental food are well intentioned and would benefit from clearer and more empirically based guidelines on how to provide foods for birds while minimizing negative consequences for wildlife (Dayer et al. 2019). The dietary needs of birds, and all wildlife, vary through seasons and life stages and thus birds need access to a variety of nutritionally rich food year-round. Perrine et al. (2025) demonstrated that, when given access to both high P:L and low P:L diets, canaries optimized their diets and had recovery times similar to those on the high P:L diet. Thus, it is possible that if wild birds are offered a variety of nutritionally rich foods, they will optimize their diet depending on their seasonal needs. To reduce the spread of disease and the risk of supplemental food spoiling, seed feeders should be frequently cleaned (∼every two weeks) and nectar feeders cleaned even more frequently (every 3-5 days) (Feliciano et al., 2018). If diseased birds are seen using the feeders, they should be taken down, cleaned, and left down for a couple weeks to reduce transmission. As human activity continues to alter the landscape of nutritional resources, wildlife in urban and suburban spaces are becoming more reliant on anthropogenic food sources. In addition to increasing risk of disease transmission via behavioral changes, reliance on anthropogenic food alters the nutritional quality of wildlife diets. Our study highlights the important role the nutritional quality of anthropogenic foods can play on host immune function and transmission and highlights the need for evidence-based recommendations and policies regarding the supplementation of food to wildlife, whether intentional or not, to protect the health of wildlife, domestic animals, and humans. Future research integrating field and experimental studies with data-driven mathematical models can provide a more robust framework for accomplishing these goals.

## Acknowledgments

We thank the Society for Comparative and Integrative Biology Divisions of Animal Behavior, Comparative Endocrinology, Ecology and Evolution, and Neurobiology, Neuroethology, and Sensory Biology for co-supporting the symposium “Cities as a Natural Experiment: How Organisms are Finding Different Solutions to the Same Urban Problems”. We also thank Lauren Eno and Wild Birds Unlimited Nature Shop in Fayetteville, AR for providing bird food sales data and Jeremy Cohen for providing the house finch photo in Figure 1. Funds were provided by grants to S.E.D. from the National Science Foundation (1941861) and Arkansas Biosciences Institute.

## References

Adamo, S. A., Bartlett, A., Le, J., Spencer, N., & Sullivan, K. (2010). Illness-induced anorexia may reduce trade-offs between digestion and immune function. Animal Behaviour, 79(1), 3–10. 10.1016/j.anbehav.2009.10.012

Adelman, J. S., Moyers, S. C., Farine, D. R., & Hawley, D. M. (2015). Feeder use predicts both acquisition and transmission of a contagious pathogen in a North American songbird. Proceedings of the Royal Society B: Biological Sciences, 282(1815), 20151429. 10.1098/rspb.2015.1429

Altizer, S., Becker, D. J., Epstein, J. H., Forbes, K. M., Gillespie, T. R., Hall, R. J., Hawley, D. M., Hernandez, S. M., Martin, L. B., Plowright, R. K., Satterfield, D. A., & Streicker, D. G. (2018). Food for contagion: Synthesis and future directions for studying host-parasite responses to resource shifts in anthropogenic environments. PHILOSOPHICAL TRANSACTIONS OF THE ROYAL SOCIETY B-BIOLOGICAL SCIENCES, 373(1745). 10.1098/rstb.2017.0102

Badyaev, A., Belloni, V., & Hill, G. (2020). House finch (Haemorhous mexicanus), version 1.0. Birds of the World.

Becker, D. J., Czirjak, G. A., Volokhov, D. V., Bentz, A. B., Carrera, J. E., Camus, M. S., Navara, K. J., Chizhikov, V. E., Fenton, M. B., Simmons, N. B., Recuenco, S. E., Gilbert, A. T., Altizer, S., & Streicker, D. G. (2018). Livestock abundance predicts vampire bat demography, immune profiles and bacterial infection risk. PHILOSOPHICAL TRANSACTIONS OF THE ROYAL SOCIETY B-BIOLOGICAL SCIENCES, 373(1745). 10.1098/rstb.2017.0089

Becker, D. J., Streicker, D. G., & Altizer, S. (2015). Linking anthropogenic resources to wildlife-pathogen dynamics: A review and meta-analysis. ECOLOGY LETTERS, 18(5), 483–495. 10.1111/ele.12428

Bonneaud, C., Tardy, L., Giraudeau, M., Hill, G. E., McGraw, K. J., & Wilson, A. J. (2019). Evolution of both host resistance and tolerance to an emerging bacterial pathogen. Evolution Letters, 3(5), 544–554. 10.1002/evl3.133

Bosse, M., Spurgin, L. G., Laine, V. N., Cole, E. F., Firth, J. A., Gienapp, P., Gosler, A. G., McMahon, K., Poissant, J., Verhagen, I., Groenen, M. A. M., Van Oers, K., Sheldon, B. C., Visser, M. E., & Slate, J. (2017). Recent natural selection causes adaptive evolution of an avian polygenic trait. Science, 358(6361), 365–368. 10.1126/science.aal3298

Brown, R. D., & Cooper, S. M. (2006). The Nutritional, Ecological, and Ethical Arguments Against Baiting and Feeding White-Tailed Deer. Wildlife Society Bulletin, 34(2), 519– 524. 10.2193/0091-7648(2006)34[519:TNEAEA]2.0.CO;2

Brunner, F. S., Schmid-Hempel, P., & Barribeau, S. M. (2014). Protein-poor diet reduces host-specific immune gene expression in *Bombus terrestris*. Proceedings of the Royal Society B: Biological Sciences, 281(1786), 20140128. 10.1098/rspb.2014.0128

Combs, M. A., Kache, P. A., VanAcker, M. C., Gregory, N., Plimpton, L. D., Tufts, D. M., Fernandez, M. P., & Diuk-Wasser, M. A. (2022). Socio-ecological drivers of multiple zoonotic hazards in highly urbanized cities. Global Change Biology, 28(5), 1705–1724. 10.1111/gcb.16033

Coogan, S. C. P., Raubenheimer, D., Zantis, S. P., & Machovsky-Capuska, G. E. (2018). Multidimensional nutritional ecology and urban birds. Ecosphere, 9(4), e02177. 10.1002/ecs2.2177

Cornelius Ruhs, E., Vezina, F., & Karasov, W. H. (2019). Physiological and Immune Responses of Free-Living Temperate Birds Provided a Gradient of Food Supplementation. PHYSIOLOGICAL AND BIOCHEMICAL ZOOLOGY, 92(1), 106–114. 10.1086/701389

Cornet, S., Bichet, C., Larcombe, S., Faivre, B., & Sorci, G. (2014). Impact of host nutritional status on infection dynamics and parasite virulence in a bird-malaria system. Journal of Animal Ecology, 83(1), 256–265. 10.1111/1365-2656.12113

Cotter, S. C., Reavey, C. E., Tummala, Y., Randall, J. L., Holdbrook, R., Ponton, F., Simpson, S. J., Smith, J. A., & Wilson, K. (2019). Diet modulates the relationship between immune gene expression and functional immune responses. Insect Biochemistry and Molecular Biology, 109, 128–141. 10.1016/j.ibmb.2019.04.009

Cox, D. T. C., & Gaston, K. J. (2016). Urban Bird Feeding: Connecting People with Nature. PLOS ONE, 11(7), e0158717. 10.1371/journal.pone.0158717

Cressler, C. E., & Adelman, J. S. (2024). Links between Innate and Adaptive Immunity Can Favor Evolutionary Persistence of Immunopathology. Integrative And Comparative Biology, 64(3), 841–852. 10.1093/icb/icae105

Cummings, C. R., Hernandez, S. M., Murray, M., Ellison, T., Adams, H. C., Cooper, R. E., Curry, S., & Navara, K. J. (2020). Effects of an anthropogenic diet on indicators of physiological challenge and immunity of white ibis nestlings raised in captivity. ECOLOGY AND EVOLUTION, 10(15), 8416–8428. 10.1002/ece3.6548

Dayer, A. A., Rosenblatt, C., Bonter, D. N., Faulkner, H., Hall, R. J., Hochachka, W. M., Phillips, T. B., & Hawley, D. M. (2019). Observations at backyard bird feeders influence the emotions and actions of people that feed birds. People and Nature, 1(2), 138–151. 10.1002/pan3.17

Dhondt, A. A., DeCoste, J. C., Ley, D. H., & Hochachka, W. M. (2014). Diverse Wild Bird Host Range of Mycoplasma gallisepticum in Eastern North America. PLOS ONE, 9(7), e103553. 10.1371/journal.pone.0103553

Dhondt, A. A., Dhondt, K. V., Hawley, D. M., & Jennelle, C. S. (2007). Experimental evidence for transmission of *Mycoplasma gallisepticum* in house finches by fomites. Avian Pathology, 36(3), 205–208. 10.1080/03079450701286277

Dhondt, A. A., Dhondt, K. V., Hochachka, W. M., Ley, D. H., & Hawley, D. M. (2017). Response of House Finches Recovered from *Mycoplasma gallisepticum* to Reinfection with a Heterologous Strain. Avian Diseases, 61(4), 437–441. 10.1637/11571-122016-Reg.1

Dhondt, A. A., States, S. L., Dhondt, K. V., & Schat, K. A. (2012). Understanding the origin of seasonal epidemics of mycoplasmal conjunctivitis. Journal of Animal Ecology, 81(5), 996–1003. 10.1111/j.1365-2656.2012.01986.x

Feliciano, L. M., Underwood, T. J., & Aruscavage, D. F. (2018). The effectiveness of bird feeder cleaning methods with and without debris. The Wilson Journal of Ornithology, 130(1), 313–320. 10.1676/16-161.1

Fleischer, R., Jones, C., Ledezma-Campos, P., Czirjak, G. A., Sommer, S., Gillespie, T. R., & Vicente-Santos, A. (2024). Gut microbial shifts in vampire bats linked to immunity due to changed diet in human disturbed landscapes. SCIENCE OF THE TOTAL ENVIRONMENT, 907. 10.1016/j.scitotenv.2023.167815

Fleming-Davies, A. E., Williams, P. D., Dhondt, A. A., Dobson, A. P., Hochachka, W. M., Leon, A. E., Ley, D. H., Osnas, E. E., & Hawley, D. M. (2018). Incomplete host immunity favors the evolution of virulence in an emergent pathogen. Science, 359(6379), 1030– 1033. 10.1126/science.aao2140

French, S. S., Webb, A. C., Wilcoxen, T. E., Iverson, J. B., DeNardo, D. F., Lewis, E. L., & Knapp, C. R. (2022). Complex tourism and season interactions contribute to disparate physiologies in an endangered rock iguana. CONSERVATION PHYSIOLOGY, 10(1). 10.1093/conphys/coac001

Hawley, D. M., Grodio, J., Frasca Jr, S., Kirkpatrick, L., & Ley, D. H. (2011). Experimental infection of domestic canaries (Serinus canaria domestica) with Mycoplasma gallisepticum: A new model system for a wildlife disease. Avian Pathology, 40(3), 321– 327.

Hawley, D. M., Thomason, C. A., Aberle, M. A., Brown, R., & Adelman, J. S. (2023). High virulence is associated with pathogen spreadability in a songbird–bacterial system. Royal Society Open Science, 10(1), 220975. 10.1098/rsos.220975

Henschen, A. E., Vinkler, M., Langager, M. M., Rowley, A. A., Dalloul, R. A., Hawley, D. M., & Adelman, J. S. (2023). Rapid adaptation to a novel pathogen through disease tolerance in a wild songbird. PLOS Pathogens, 19(6), e1011408. 10.1371/journal.ppat.1011408

Hite, J. L., & Cressler, C. E. (2019). Parasite-Mediated Anorexia and Nutrition Modulate Virulence Evolution. Integrative and Comparative Biology, 59(5), 1264–1274. 10.1093/icb/icz100

Hite, J. L., Pfenning, A. C., & Cressler, C. E. (2020). Starving the Enemy? Feeding Behavior Shapes Host-Parasite Interactions. Trends in Ecology & Evolution, 35(1), 68–80. 10.1016/j.tree.2019.08.004

Hochachka, W. M., & Dhondt, A. A. (2000). Density-dependent decline of host abundance resulting from a new infectious disease. Proceedings of the National Academy of Sciences, 97(10), 5303–5306.

Hochachka, W. M., Dobson, A. P., Hawley, D. M., & Dhondt, A. A. (2021). Host population dynamics in the face of an evolving pathogen. Journal of Animal Ecology, 1365-2656.13469. 10.1111/1365-2656.13469

Horn, D. J., Johansen, S. M., & Wilcoxen, T. E. (2014). Seed and feeder use by birds in the United States and Canada. Wildlife Society Bulletin, 38(1), 18–25. 10.1002/wsb.365

Hosseini, P. R., Dhondt, A. A., & Dobson, A. (2004). Seasonality and wildlife disease: How seasonal birth, aggregation and variation in immunity affect the dynamics of *Mycoplasma gallisepticum* in house finches. Proceedings of the Royal Society of London. Series B: Biological Sciences, 271(1557), 2569–2577. 10.1098/rspb.2004.2938

Hwang, J., Kim, Y., Lee, S.-W., Kim, N.-Y., Chun, M.-S., Lee, H., & Gottdenker, N. (2018). Anthropogenic food provisioning and immune phenotype: Association among supplemental food, body condition, and immunological parameters in urban environments. ECOLOGY AND EVOLUTION, 8(5), 3037–3046. 10.1002/ece3.3814

Ingala, M. R., Becker, D. J., Holm, J. B., Kristiansen, K., & Simmons, N. B. (2019). Habitat fragmentation is associated with dietary shifts and microbiota variability in common vampire bats. ECOLOGY AND EVOLUTION, 9(11), 6508–6523. 10.1002/ece3.5228

Jahanian, R. (2009). Immunological responses as affected by dietary protein and arginine concentrations in starting broiler chicks. Poultry Science, 88(9), 1818–1824. 10.3382/ps.2008-00386

Johansen, S. M., Horn, D. J., & Wilcoxen, T. E. (2014). Factors Influencing Seed Species Selection by Wild Birds at Feeders. The Wilson Journal of Ornithology, 126(2), 374–381. 10.1676/13-066.1

Jones, J. D., Kauffman, M. J., Monteith, K. L., Scurlock, B. M., Albeke, S. E., & Cross, P. C. (2014). Supplemental feeding alters migration of a temperate ungulate. Ecological Applications, 24(7), 1769–1779. 10.1890/13-2092.1

Knutie, S. A. (2020). Food supplementation affects gut microbiota and immunological resistance to parasites in a wild bird species. Journal of Applied Ecology, 57(3), 536–547. 10.1111/1365-2664.13567

Kollias, G. V., Sydenstricker, K. V., Kollias, H. W., Ley, D. H., Hosseini, P. R., Connolly, V., & Dhondt, A. A. (2004). Experimental infection of House Finches with Mycoplasma gallisepticum. Journal of Wildlife Diseases, 40(1), 79–86. 10.7589/0090-3558-40.1.79

Ley, D. H., & Yoder Jr, H. (2008). Mycoplasma gallisepticum infection. Diseases of Poultry, 12, 807–834.

Li, M. Y., & Muldowney, J. S. (1995). Global stability for the SEIR model in epidemiology. Mathematical Biosciences, 125(2), 155–164.

Lochmiller, R. L., Vestey, M. R., & Boren, J. C. (1993). Relationship between protein nutritional status and immunocompetence in northern bobwhite chicks. The Auk, 110(3), 503–510.

Love, A. C., Tabb, V., Youssef, N. H., Wilder, S. M., & DuRant, S. E. (2024). Effect of dietary macronutrients and immune challenge on gut microbiota, physiology and feeding behaviour in zebra finches. Molecular Ecology, 33(14), e17428. 10.1111/mec.17428

Moyers, S. C., Adelman, J. S., Farine, D. R., Thomason, C. A., & Hawley, D. M. (2018). Feeder density enhances house finch disease transmission in experimental epidemics. Philosophical Transactions of the Royal Society B: Biological Sciences, 373(1745), 20170090. 10.1098/rstb.2017.0090

Murray, M. H., Becker, D. J., Hall, R. J., & Hernandez, S. M. (2016). Wildlife health and supplemental feeding: A review and management recommendations. BIOLOGICAL CONSERVATION, 204(B), 163–174. 10.1016/j.biocon.2016.10.034

Murray, M. H., Cembrowski, A., Latham, A. D. M., Lukasik, V. M., Pruss, S., & St Clair, C. C. (2015). Greater consumption of protein-poor anthropogenic food by urban relative to rural coyotes increases diet breadth and potential for human–wildlife conflict. Ecography, 38(12), 1235–1242. 10.1111/ecog.01128

Murray, M. H., Kidd, A. D., Curry, S. E., Hepinstall-Cymerman, J., Yabsley, M. J., Adams, H. C., Ellison, T., Welch, C. N., & Hernandez, S. M. (2018). From wetland specialist to hand-fed generalist: Shifts in diet and condition with provisioning for a recently urbanized wading bird. Philosophical Transactions of the Royal Society B: Biological Sciences, 373(1745), 20170100. 10.1098/rstb.2017.0100

Murray, M. H., Sánchez, C. A., Becker, D. J., Byers, K. A., Worsley-Tonks, K. E., & Craft, M. E. (2019). City sicker? A meta-analysis of wildlife health and urbanization. Frontiers in Ecology and the Environment, 17(10), 575–583. 10.1002/fee.2126

Orros, M. E., & Fellowes, M. D. E. (2012). Supplementary feeding of wild birds indirectly affects the local abundance of arthropod prey. Basic and Applied Ecology, 13(3), 286– 293. 10.1016/j.baae.2012.03.001

Perrine, W. G., Sauer, E. L., Love, A. C., Morris, A., Novotny, J., & DuRant, S. E. (2025). A high lipid diet leads to greater pathology and lower tolerance during infection. Journal of Experimental Biology, jeb.249541. 10.1242/jeb.249541

Pike, V. L., Lythgoe, K. A., & King, K. C. (2019). On the diverse and opposing effects of nutrition on pathogen virulence. Proceedings of the Royal Society B: Biological Sciences, 286(1906), 20191220. 10.1098/rspb.2019.1220

Plummer, K. E., Bearhop, S., Leech, D. I., Chamberlain, D. E., & Blount, J. D. (2013). Winter food provisioning reduces future breeding performance in a wild bird. Scientific Reports, 3(1), 2002. 10.1038/srep02002

Ponton, F., Morimoto, J., Robinson, K., Kumar, S. S., Cotter, S. C., Wilson, K., & Simpson, S. J. (2020). Macronutrients modulate survival to infection and immunity in *Drosophila*. Journal of Animal Ecology, 89(2), 460–470. 10.1111/1365-2656.13126

Povey, S., Cotter, S. C., Simpson, S. J., & Wilson, K. (2014). Dynamics of macronutrient self-medication and illness-induced anorexia in virally infected insects. Journal of Animal Ecology, 83(1), 245–255. 10.1111/1365-2656.12127

Robb, G. N., McDonald, R. A., Chamberlain, D. E., & Bearhop, S. (2008). Food for thought: Supplementary feeding as a driver of ecological change in avian populations. Frontiers in Ecology and the Environment, 6(9), 476–484. 10.1890/060152

RStudio Team. (2021). RStudio: Integrated Development Environment for R. RStudio, PBC. http://www.rstudio.com/

Ruden, R. M., & Adelman, J. S. (2021). Disease tolerance alters host competence in a wild songbird. Biology Letters, 17(10), 20210362. 10.1098/rsbl.2021.0362

Sauer, E. L., Connelly, C., Perrine, W., Love, A. C., & DuRant, S. E. (2023). Male pathology regardless of behaviour drives transmission in an avian host–pathogen system. Journal of Animal Ecology, 93(1), 36–44. 10.1111/1365-2656.14026

Sauer, E. L., Stacy, C., Perrine, W., Love, A. C., Lewis, J. A., & DuRant, S. E. (2024). Diet driven differences in host tolerance are linked to shifts in global gene expression in a common avian host-pathogen system. 10.1101/2024.08.07.607042

Schell, C. J., Stanton, L. A., Young, J. K., Angeloni, L. M., Lambert, J. E., Breck, S. W., & Murray, M. H. (2021). The evolutionary consequences of human–wildlife conflict in cities. Evolutionary Applications, 14(1), 178–197. 10.1111/eva.13131

Semeniuk, C. A. D., Bourgeon, S., Smith, S. L., & Rothley, K. D. (2009). Hematological differences between stingrays at tourist and non-visited sites suggest physiological costs of wildlife tourism. Biological Conservation, 142(8), 1818–1829. 10.1016/j.biocon.2009.03.022

Semeniuk, C., & Rothley, K. (2008). Costs of group-living for a normally solitary forager: Effects of provisioning tourism on southern stingrays Dasyatis americana. Marine Ecology Progress Series, 357, 271–282. 10.3354/meps07299

Shutt, J. D., & Lees, A. C. (2021). Killing with kindness: Does widespread generalised provisioning of wildlife help or hinder biodiversity conservation efforts? Biological Conservation, 261, 109295. 10.1016/j.biocon.2021.109295

Soetaert, K., Petzoldt, T., & Setzer, R. W. (2010). Package deSolve: Solving initial value differential equations in R. Journal of Statistical Software, 33(9), 1–25.

Stewart, K., Norton, T., Mohammed, H., Browne, D., Clements, K., Thomas, K., Yaw, T., & Horrocks, J. (2016). EFFECTS OF “SWIM WITH THE TURTLES” TOURIST ATTRACTIONS ON GREEN SEA TURTLE (*CHELONIA MYDAS*) HEALTH IN BARBADOS, WEST INDIES. JOURNAL OF WILDLIFE DISEASES, 52(2), S104–S117. 10.7589/52.2S.S104

Strandberg, L., Verdrengh, M., Enge, M., Andersson, N., Amu, S., Önnheim, K., Benrick, A., Brisslert, M., Bylund, J., Bokarewa, M., Nilsson, S., & Jansson, J.-O. (2009). Mice Chronically Fed High-Fat Diet Have Increased Mortality and Disturbed Immune Response in Sepsis. PLoS ONE, 4(10), e7605. 10.1371/journal.pone.0007605

Strandin, T., Babayan, S. A., & Forbes, K. M. (2018). Reviewing the effects of food provisioning on wildlife immunity. Philosophical Transactions of the Royal Society B: Biological Sciences, 373(1745), 20170088. 10.1098/rstb.2017.0088

Sudnick, M. C., DuRant, S. E., & Sauer, E. L. (2025). Prior infection induces long-lasting partial immunity to reduce transmission within flocks in an avian host-pathogen system. 10.1101/2025.03.23.644804

Taylor, A. K., Cao, W., Vora, K. P., Cruz, J. D. L., Shieh, W.-J., Zaki, S. R., Katz, J. M., Sambhara, S., & Gangappa, S. (2013). Protein Energy Malnutrition Decreases Immunity and Increases Susceptibility to Influenza Infection in Mice. The Journal of Infectious Diseases, 207(3), 501–510. 10.1093/infdis/jis527

Viquez-R, L., Henrich, M., Riegel, V., Bader, M., Wilhelm, K., Heurich, M., & Sommer, S. (2024). A taste of wilderness: Supplementary feeding of red deer (*Cervus elaphus*) increases individual bacterial microbiota diversity but lowers abundance of important gut symbionts. ANIMAL MICROBIOME, 6(1). 10.1186/s42523-024-00315-6

Wilcoxen, T. E., Horn, D. J., Hogan, B. M., Hubble, C. N., Huber, S. J., Flamm, J., Knott, M., Lundstrom, L., Salik, F., Wassenhove, S. J., & Wrobel, E. R. (2015). Effects of bird-feeding activities on the health of wild birds. Conservation Physiology, 3(1), cov058. 10.1093/conphys/cov058

Yadav, J. P., Tomar, P., Singh, Y., & Khurana, S. K. (2022). Insights on *Mycoplasma gallisepticum* and *Mycoplasma synoviae* infection in poultry: A systematic review. Animal Biotechnology, 33(7), 1711–1720. 10.1080/10495398.2021.1908316

